## Supplementary Information for "Genome-wide effects of the antimicrobial peptide apidaecin on translation termination"

### SUPPLEMENTARY FIGURES AND TABLES

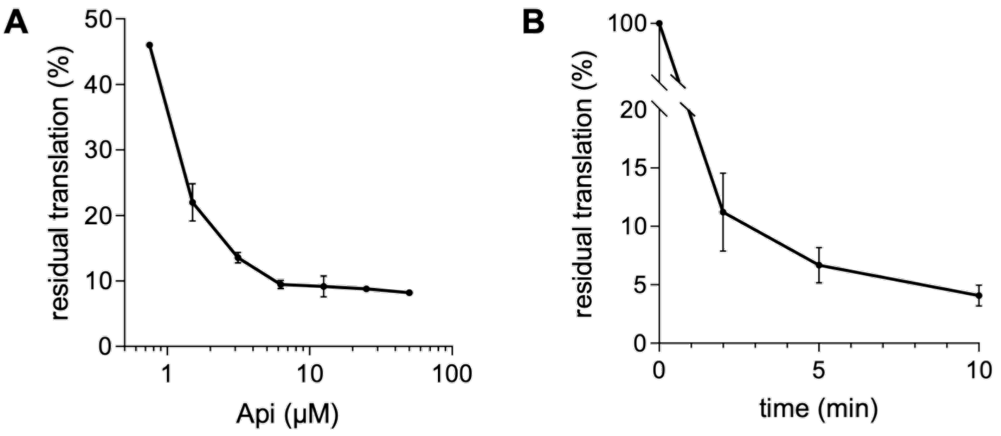


**Associated to Figure 1**

**Figure supplement 1.** Api inhibits global protein synthesis. (**A**) Residual protein synthesis in *E. coli* cells (strain BL21 Δ*tolC*) exposed for 1 min to varying concentrations of Api137, as estimated by L-[$^{35}$S]-methionine incorporation into TCA-precipitable proteins. Incorporation of L-[$^{35}$S]-methionine in a sample devoid of Api was set as 100%. (**B**) Time course of inhibition of protein synthesis in cells exposed to 6.25 μM (4X MIC) of Api137. The shown data is the average of two independent experiments. Error bars indicate s.e.m.

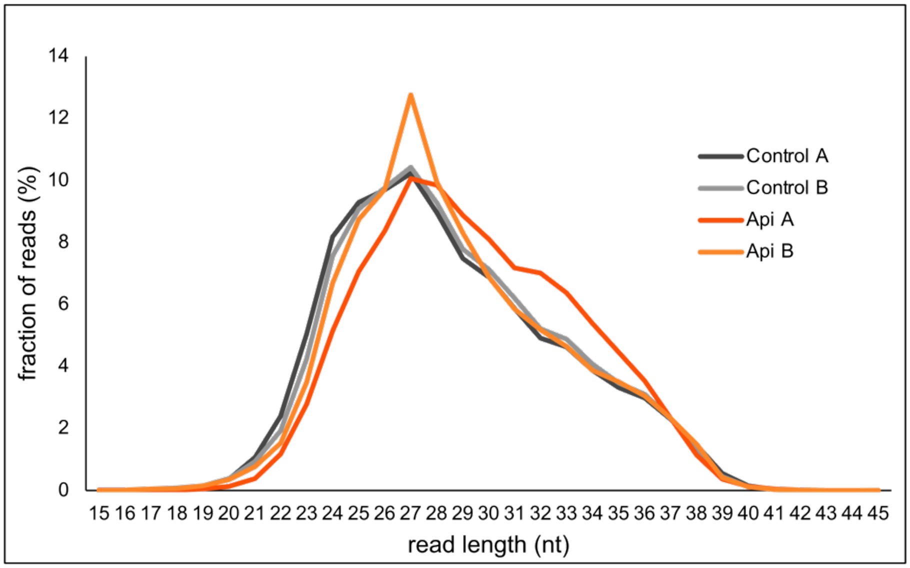


**Associated to Figure 2**

**Figure supplement 1.** Footprint length distributions for Api and control samples. Length distribution of ribosome footprint reads aligned to the genome after filtering out non-coding RNA reads.

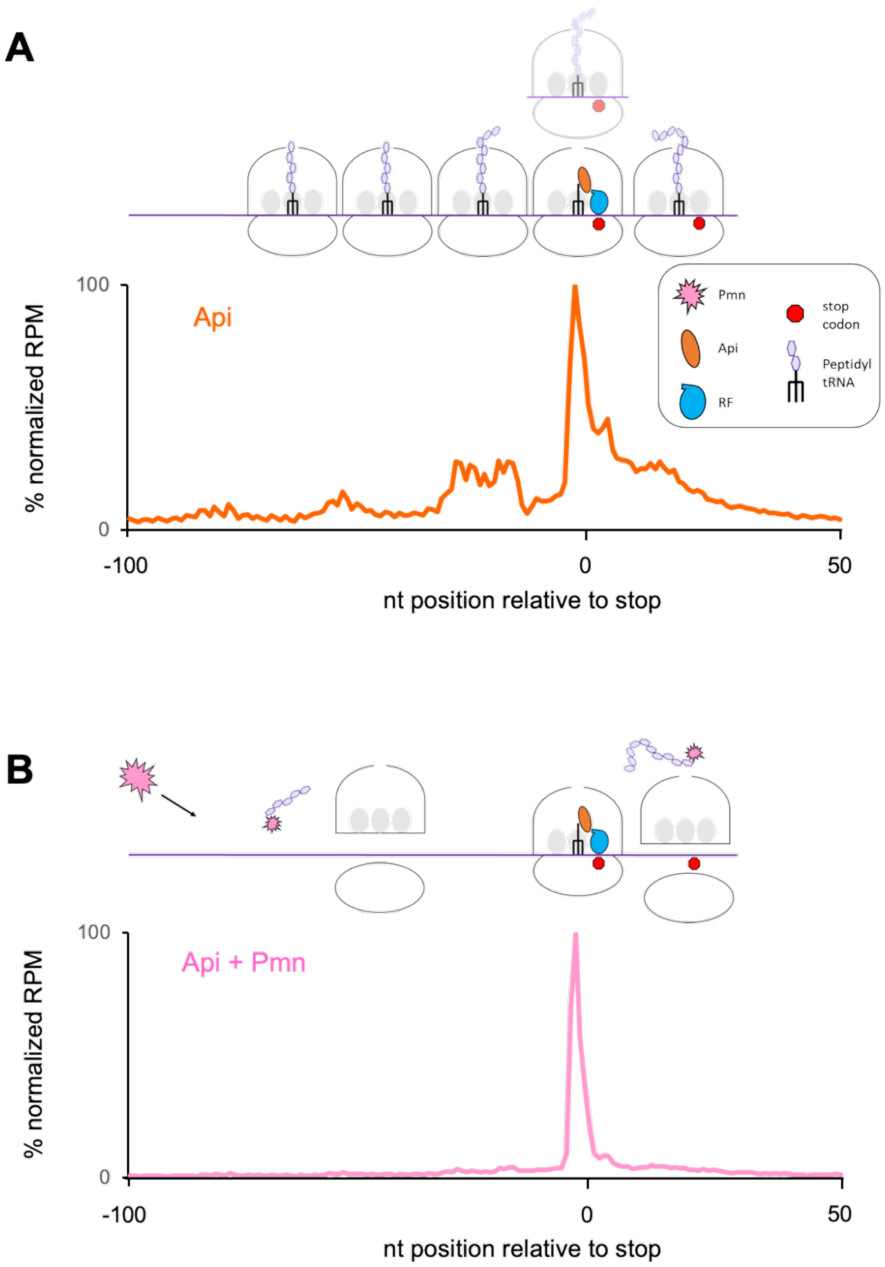


**Associated to Figure 2**

**Figure supplement 2.** Puromycin treatment simplifies the ribosome distribution pattern in Api-treated cells. Metagene plots of average normalized ribosome density in the vicinity of stop codons of genes in cells exposed to Api without (*top*) or with (*bottom*) puromycin (Pmn) treatment. Pmn treatment removes peptidyl-tRNA and facilitates dissociation of ribosomes from mRNA. Ribosomes arrested at the stop codon in the post-release state are expected to be refractory to Pmn action. Note that the metagene plots show the relative (normalized) distribution of ribosomes rather than the absolute number of ribosomes associated with each mRNA nucleotide.

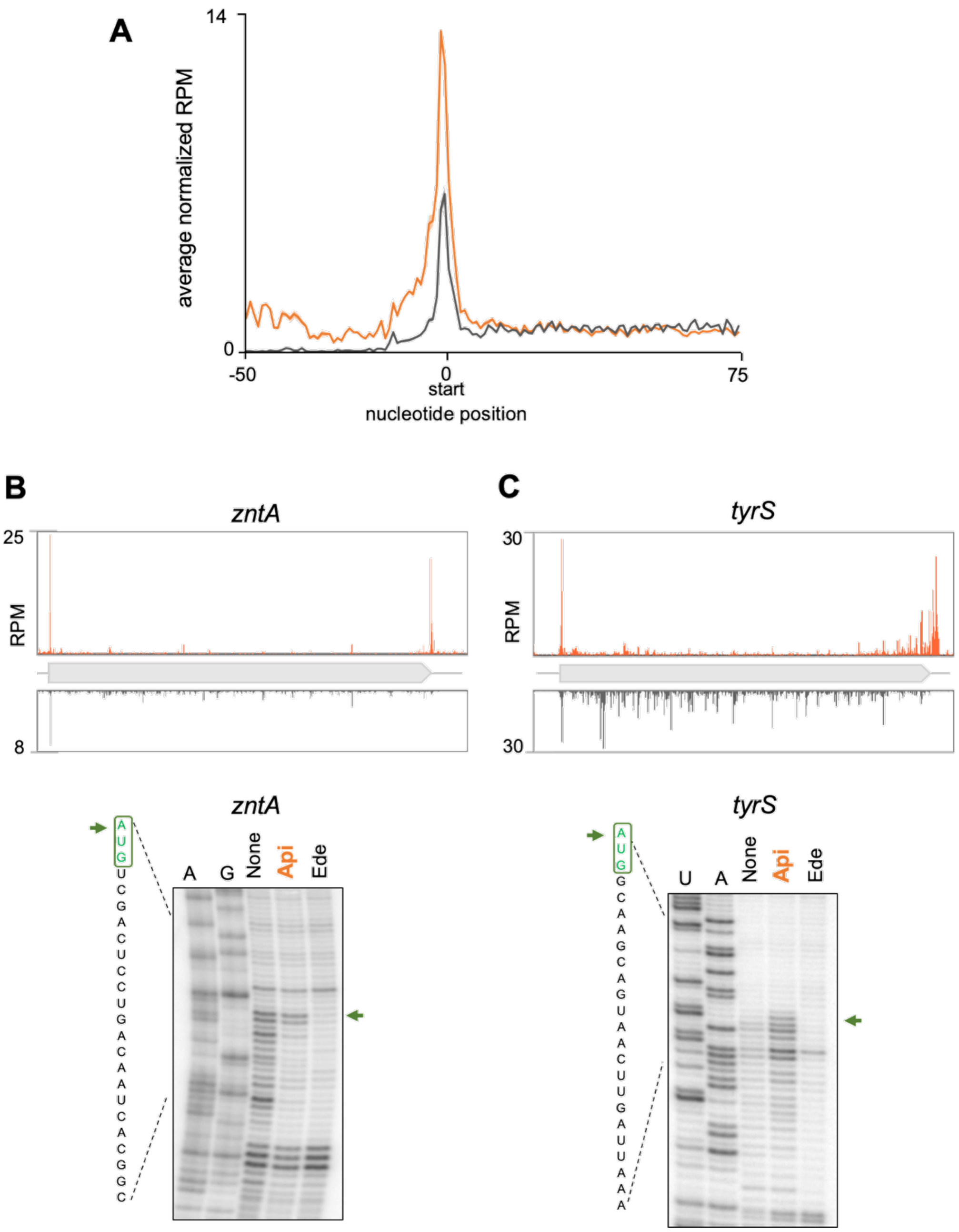


**Associated to Figure 2**

**Figure supplement 3.** Api moderately increases ribosome density at start codons. (**A**) Metagene plot of the average normalized ribosome density in the vicinity of the start codons in *E. coli* cells treated (orange) or not (gray) with Api. (**B**) and (**C**) Examples of genes where Api causes ribosome stalling at the start codons during in vivo and in vitro translation. *Top*: Ribosome footprint density observed in Ribo-seq experiments. *Bottom*: toeprinting analysis showing ribosome stalling at the start codons (green arrows) during cell-free translation in the presence of 2 mM of Api. The control antibiotic edein (Ede), which prevents ribosome binding to mRNA, was used to distinguish the toeprint bands originating from translating ribosomes.

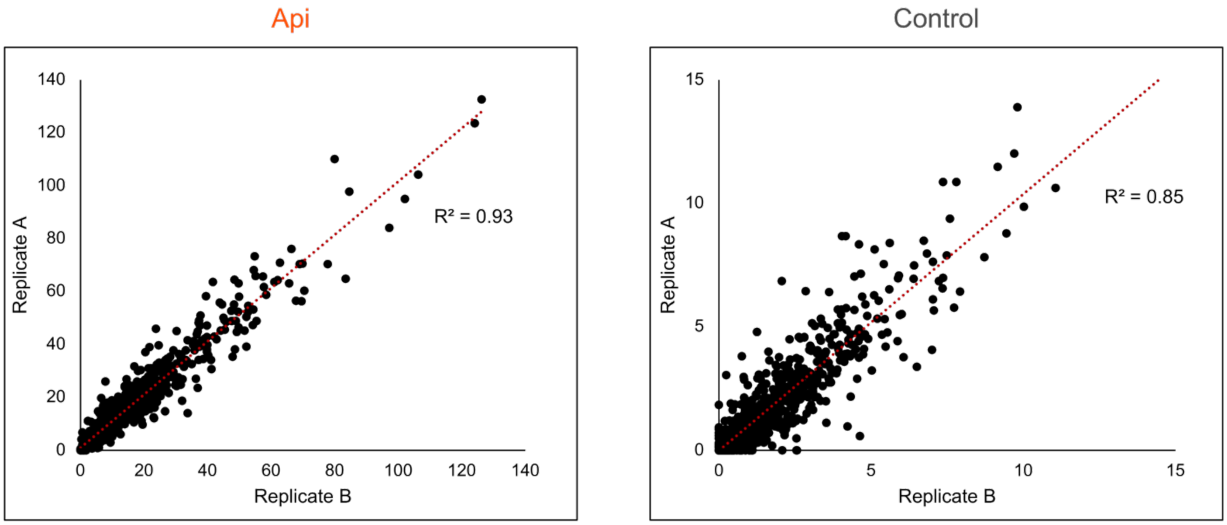


**Associated to Figure 3**

**Figure supplement 1.** Reproducibility of stop codon effects. Correlation of the stop codon (SC) scores in Api-treated or untreated (control) cells estimated from the Ribo-seq data obtained in two independent biological replicates.

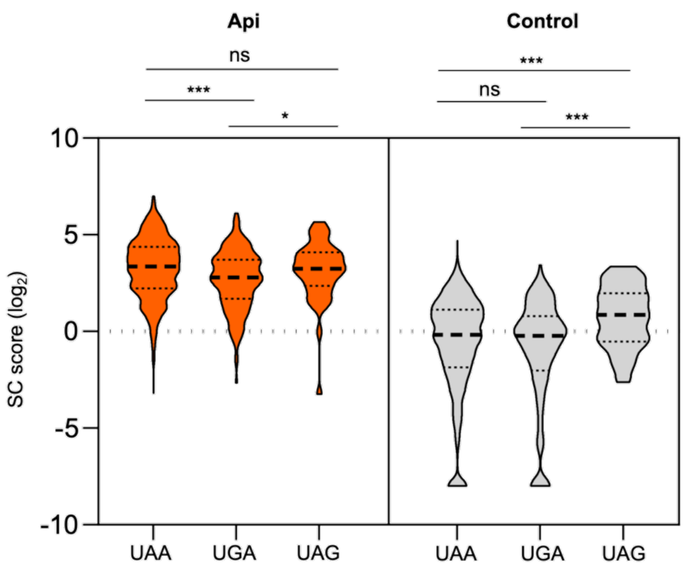


**Associated to Figure 3**

**Figure supplement 2.** Stop codon scores at different stop codons. Violin plots of genome-wide Stop codon (SC) scores binned by stop codon identity. Two-sided Mann-Whitney *U* tests were performed to assess the significance of difference between genes grouped by stop codons: Api UAA vs UGA $P$ = 1.2E-5, Api UGA vs UAG $P$ = 0.032, control UAA vs UAG $P$ = 5.2E-4, control UGA vs UAG $P$ = 1.5E-4.

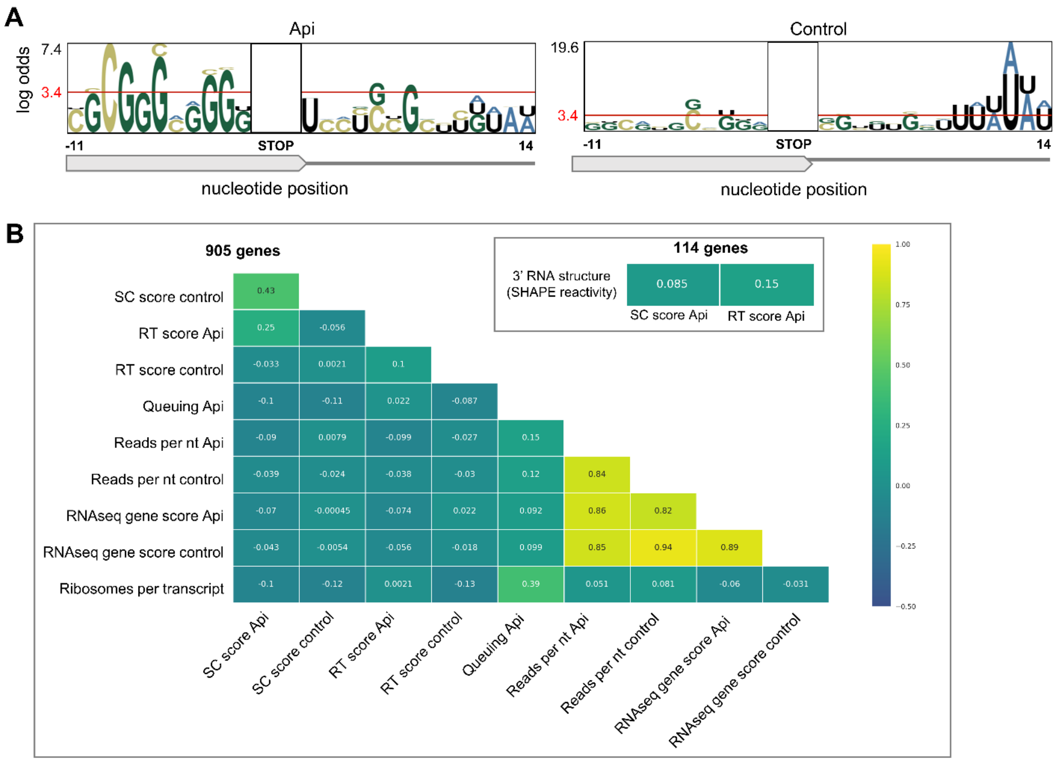


#### Associated to Figure 3

**Figure supplement 3.** Evaluation of cellular factors that could potentially contribute to Api-mediated ribosome arrest at stop codons. (**A**) pLogo analysis of nucleotide sequences in the vicinity of the termination sites of the 50% high-SC score vs. 50% low SC-score genes in Api treated (*left*) and control (*right*) cells. (**B**) Correlation between SC scores and readthrough (RT) scores of genes from Api treated cells with various parameters calculated for the 905 well-separated (50 nt away from the nearest gene) and actively translated (ribosome footprint (rfp) density per nt ≥0.1) genes. Other than SC score and RT score, the following parameters were used: Asymmetry score: a parameter reflecting difference in the number of rfps in the first half of the gene relative to the number of rfps in the second half of the gene (Mohammad et al., 2019); rfp density: the total number of rfps within an ORF (excluding the first and last 9 nts) divided by the length of an ORF; RNA-seq gene score: total number of reads per million mapped to an ORF; rfp per transcript: the number of rfp mapped to an ORF (excluding the first and last 9 nts) divided by the RNA-seq gene score (formally similar to translation efficiency metrics which is poorly applicable for the Api sample); 3' mRNA structure: the mean SHAPE reactivity of mRNA segment that includes 40 nt upstream and 10 nts downstream of the stop codon (Huang et al., 2019). The latter was calculated for 114 out of the aforementioned 905 genes, for which SHAPE-seq data were available.

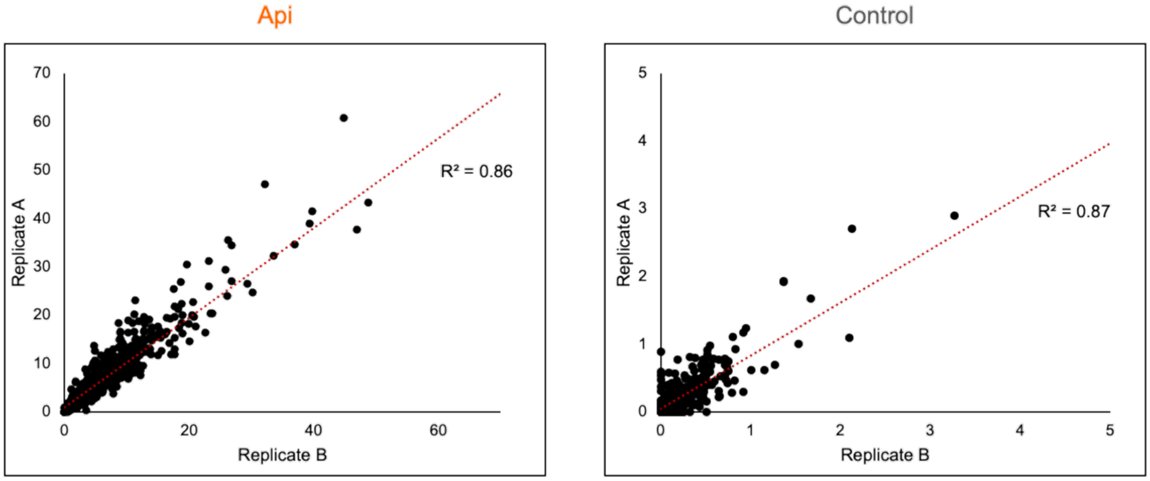


**Associated to Figure 4**

**Figure supplement 1.** Reproducibility of stop codon bypass effects. Correlation of the readthrough (RT) scores in Api-treated or control cells estimated from the Ribo-seq data obtained in two independent biological replicates.

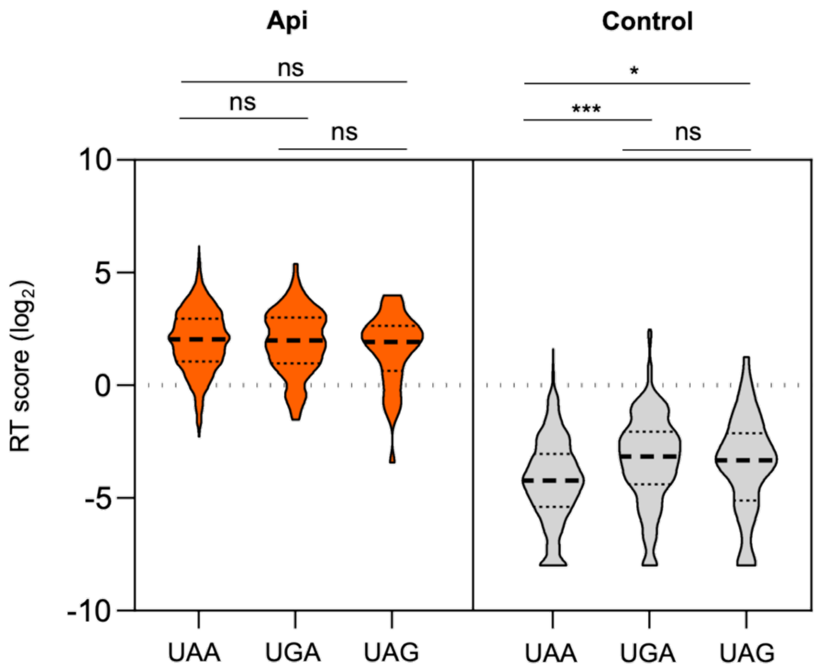


**Associated to Figure 4**

**Figure supplement 2.** Readthrough scores at different stop codons. Violin plots of genome-wide Readthrough (RT) scores binned by stop codon identity. Two-sided Mann-Whitney *U* tests were performed to assess the significance of difference between genes grouped by stop codons: Control UAA vs UGA $P$ = 5.91E-9, control UAA vs UAG $P$ = 0.012.

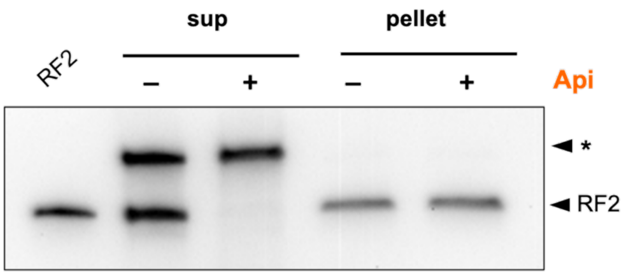


**Associated to Figure 4**

**Figure supplement 3.** Api treatment depletes free RF2 in the cell. Lysates of the untreated or Api-treated cells were layered over sucrose cushion and centrifuged for 1 hour at 100,000 rpm in a TLA-100.2 rotor at 4 °C. The presence of RF2 in the ribosome pellet or in the post-ribosomal supernatant was tested by Western blotting using anti-RF2 serum. The purified *E. coli* $His_6$-tagged RF2 was used as a control (lane marked ‘RF2’). 10 times the protein amount was loaded onto the lanes with supernatant samples in comparison with the pellet samples. The unknown protein cross-reacting with the anti-RF2 serum (indicated with an asterisks) serves as a loading control.

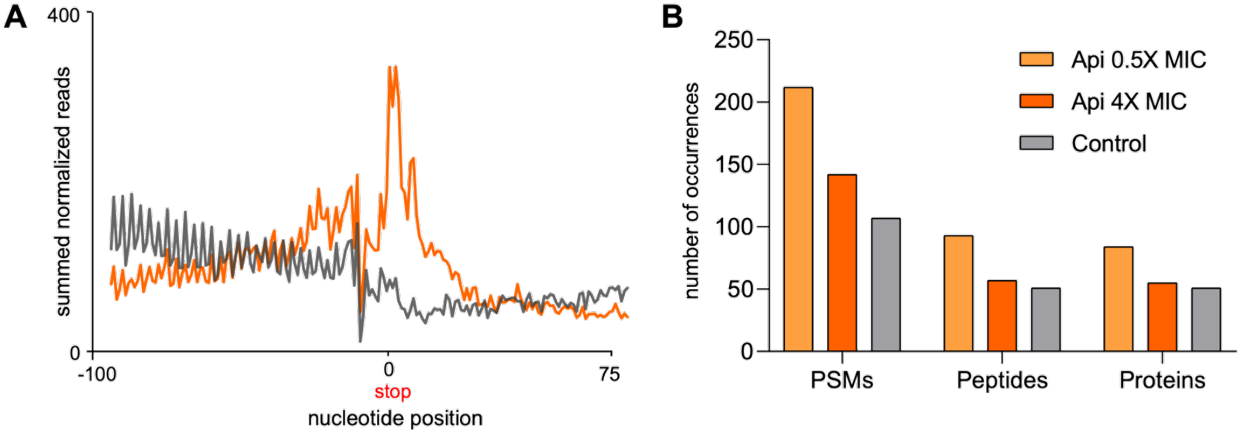


**Associated to Figure 4**

**Figure supplement 4.** In the Api-treated cells, ribosomes translate mRNA segments downstream from the stop codon of the main ORF. (**A**) Metagene plot of the ribosome footprints density in the vicinity of the first in-frame stop codon downstream from the main ORF termination codon in cells treated (orange) or not (gray) with Api. (**B**) Number of proteins with C-terminal extensions or PSMs and peptides counts belonging to the C-terminal extensions of proteins in the control and Api-treated cells identified by shotgun proteomics (a more relaxed criteria compared to those used for generation of the Figure 4B plots, see Materials and Methods).

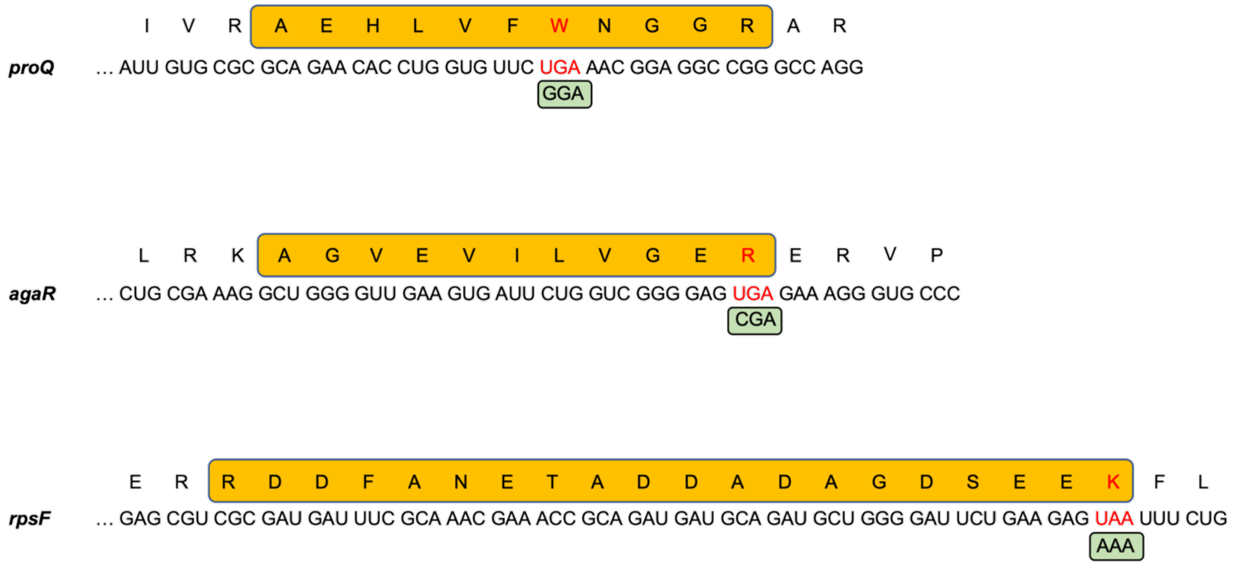


**Associated to Figure 4**

**Figure supplement 5.** Stop codon bypass by near-cognate aminoacyl-tRNA misincorporation. Api-induced 0-frame stop codon bypass occurs via decoding of the stop codon by a near-cognate aminoacyl-tRNA**.** Three peptides (yellow boxes), encoded by the mRNA sequences spanning the stop codons were identify by mass-spectrometry in the cells exposed to 0.5x MIC of Api. The nucleotide sequences of the 3'-proximal segments of the corresponding ORFs and amino acid sequences of the encoded proteins are shown. Stop codons and amino acids erroneously incorporated at stop codons are shown in red. The codon recognized by aminoacyl-tRNA that likely decoded the stop codon is shown in green boxes.

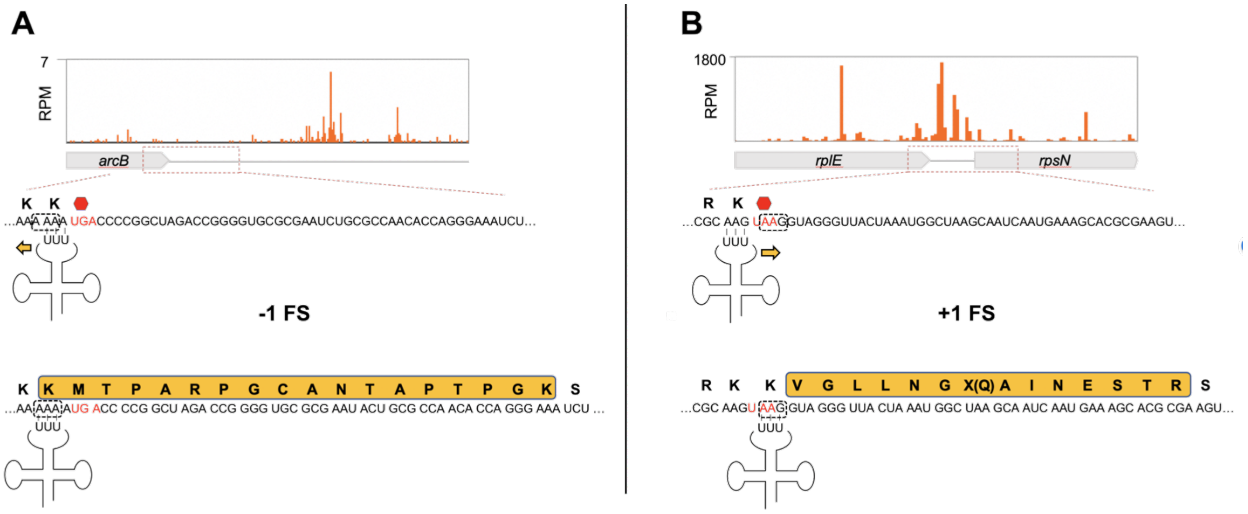


**Associated to Figure 4**

**Figure supplement 6.** Possible scenarios for stop codon bypass via frameshifting in Api-treated cells**. (A)** *Top*: Ribosome footprints density downstream of the *arcB* ORF in Api-treated cells. *Bottom*: Inability of the ribosome to rapidly terminate translation at the *arcB* UGA stop codon (red), due to Api-induced RF2 depletion, may allow for to a backwards shift of the peptidyl-tRNA$^{Lys}$ occupying the last *arcB* sense codon AAA (boxed) within the slippery sequence, landing in an identical (-1) frame codon (boxed). (**B**) *Top*: Ribosome footprints density in the region downstream of the *rplE* ORF in Api-treated cells. *Bottom*: Slow termination of translation at the *rplE* UAA stop codon (red), may lead to repairing of the peptidyl-tRNA$^{Lys}$ occupying the last *rplE* sense codon AAG to the nearby identical codon in a (+1) frame (boxed). In (A) and (B), the amino acid sequences of the tryptic peptide identified by shotgun proteomics in cells treated with Api are highlighted in yellow. Note that the stop codon within the (+1) frame downstream of *rplE* is apparently bypassed by misincorporation of a near-cognate Gln-tRNA$^{(UUG)}$. Also of note, the same tryptic peptide as the one encoded in the mRNA segment downstream of the *arcB* gene could be also identified in untreated cells.

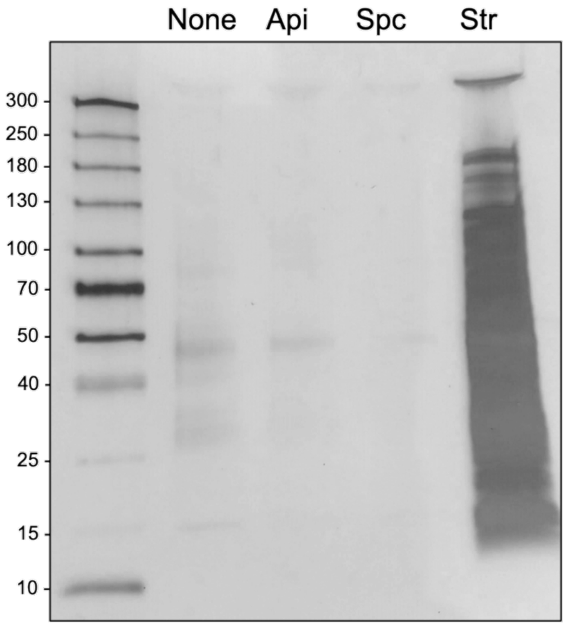


**Associated to Figure 4**

**Figure supplement 7.** Api treatment does not lead to pronounced protein aggregation in *E. coli* cells. Cells were treated or not with apidaecin (Api), spectinomycin (Spc), or streptomycin (Str) at 2-fold the respective MICs for 30 minutes. Protein aggregates were isolated, separated by SDS-PAGE and visualized by silver staining. Spc and Str are verified positive and negative controls in these conditions (Ling et al., 2012).

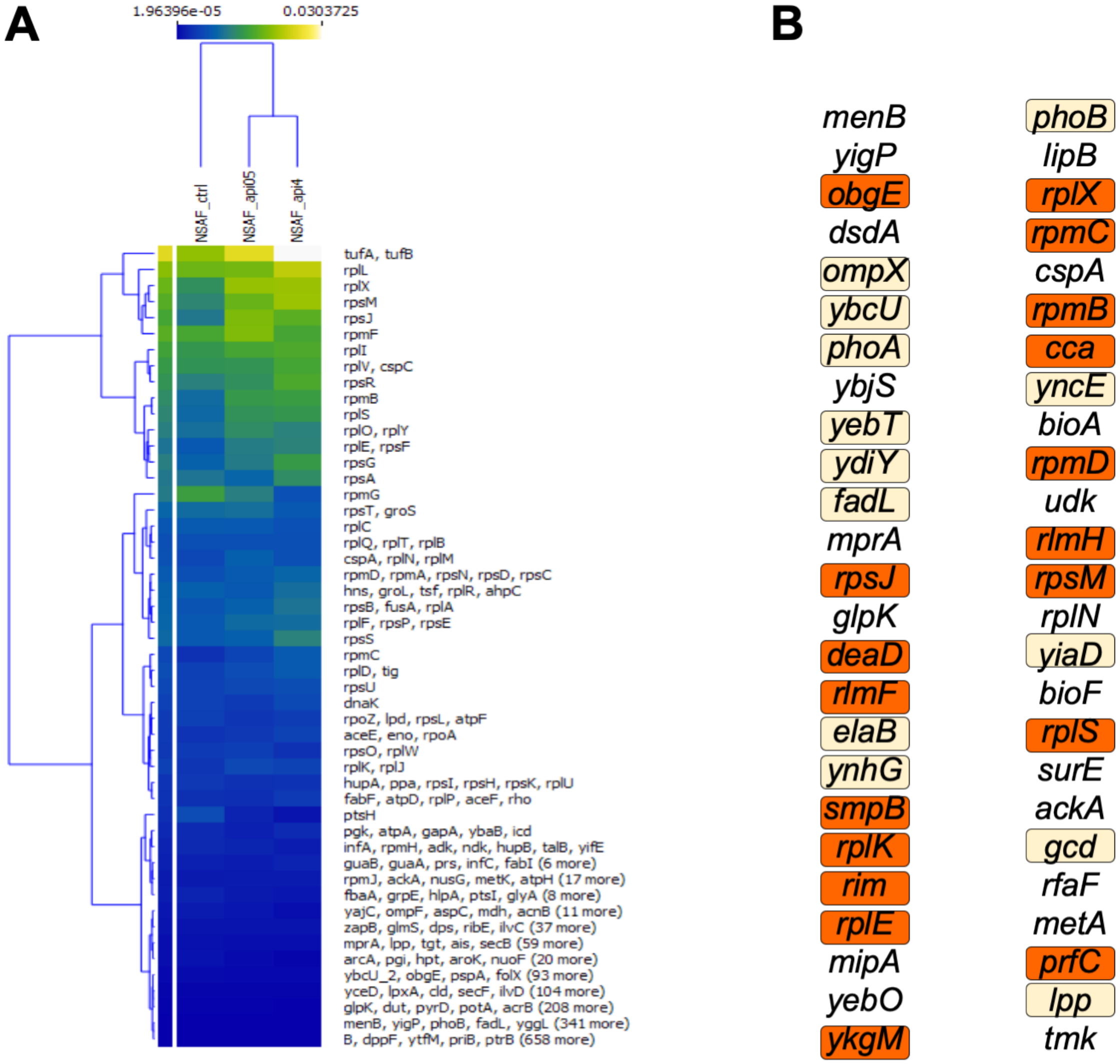


**Associated to Figure 5**

**Figure supplement 1.** Api induced changes in protein abundance. **(A)** Api-induced changes in protein abundance. Changes in relative protein abundance in *E. coli* cells exposed to 0.5x MIC and 4x MIC of Api relative to the untreated control cells. The heatmap scale reflects length-adjusted fractional abundance of the protein in the total protein sample (blue – less abundant, yellow – more abundant). Genes are clustered using hierarchical k-mean clustering algorithm. The color represents the relative abundance of each protein or protein cluster using normalized spectral abundance factor (NSAF). **(B)** Top 50 proteins showing significant increase in abundance in the cells treated with 0.5x MIC of Api relative to the untreated control. Proteins related to translation are highlighted in orange. Periplasmic, inner- and outer-membrane proteins are highlighted in beige.

Minimal inhibitory concentration

| *E. coli* strain | Api MIC (μM) |
|---|---|
| BW25113 (wt) | 12.5 |
| BW25113 Δ*arfA* | 6.25 |
| BW25113 Δ*arfB* | 12.5 |
| BW25113 Δ*smpB* | 25 |

**Associated to Figure 5**

**Figure supplement 2.** Sensitivity towards Api of cells lacking ribosome rescue systems. MIC values of Api against *E. coli* BW25113 cells lacking individual ribosome rescue systems. The SmpB protein is essential for the operation of the tmRNA-based ribosome rescue system (Buskirk and Green, 2017). All the tested strains were acquired from the Keio collection (Baba et al., 2006) and their identity was verified by PCR. Note that a two-fold difference in MIC is considered to be within the experimental error.

**Supplementary Information references**


Baba, T., Ara, T., Hasegawa, M., Takai, Y., Okumura, Y., Baba, M., Datsenko, K.A., Tomita, M., Wanner, B.L., and Mori, H. (2006). Construction of *Escherichia coli* K-12 in-frame, single-gene knockout mutants: the Keio collection. Mol. Syst. Biol. *2*, 2006 0008.

Buskirk, A.R., and Green, R. (2017). Ribosome pausing, arrest and rescue in bacteria and eukaryotes. Philos. Trans. R. Soc. Lond. B Biol. Sci *372*, 1716

Huang, L., Aghajan, M., Quesenberry, T., Low, A., Murray, S.F., Monia, B.P., and Guo, S. (2019). Targeting translation termination machinery with antisense oligonucleotides for diseases caused by nonsense mutations. Nucleic Acid Ther., *29*, 175-186.

Ling, J., Cho, C., Guo, L.T., Aerni, H.R., Rinehart, J., and Soll, D. (2012). Protein aggregation caused by aminoglycoside action is prevented by a hydrogen peroxide scavenger. Mol. Cell *48*, 713-722.

Mohammad, F., Green, R., and Buskirk, A.R. (2019). A systematically-revised ribosome profiling method for bacteria reveals pauses at single-codon resolution. eLife *8*, e42591
